# ENSO-driven temperature variability shifts the altitudinal frontier of dengue risk in the Andes

**DOI:** 10.64898/2026.03.06.710067

**Authors:** Adrià San-José, Daniela Puentes, Rachel Lowe, Mauricio Santos-Vega

## Abstract

Despite the high burden of dengue in Latin America, high elevations have historically protected megacities such as Mexico City, Quito, and Bogotá. Climate change is expected to shift elevational suitability frontiers, yet the mechanisms governing disease expansion into highland regions and the timescales over which it occurs remain poorly characterized. Using spatiotemporal decomposition techniques, we identified a mechanistic cascade linking (1) the El Niño–Southern Oscillation (ENSO) to local Colombian climate, accounting for ∼85% of interannual temperature variability and ∼53% of rainfall variability, and (2) local climate to dengue dynamics, accounting for ∼66–76% of variation in incidence and ∼44–55% of its altitudinal range. Multiple lines of evidence, including spatial ENSO signatures across Andean ridges, temporal predictive performance, and vectorial-capacity modeling, identified the local temperature expression of the ENSO teleconnection as the dominant driver of transmission. During warm ENSO phases, dengue incidence increases exponentially and transmission expands into higher elevations. This shifts the altitudinal distribution of dengue transmission upward, exposing immunologically naïve populations in highland areas that were previously considered unsuitable. Mechanistically, this altitudinal shift arises because a coherent warming across elevational strata disproportionately increases transmission in highland regions, where small temperature increments produce the largest increase in vectorial capacity. These findings challenge the view that disease expansion is primarily a gradual response to long-term warming and highlight the importance of climate variability for disease projection, attribution, and public health preparedness.

**Significance statement:** While increasing mean temperatures are a known threat to the changing landscape of infectious diseases, this study reveals that interannual climate variability can be a more immediate driver of disease burden and expansion. We show that the El Niño phenomenon produces local temperature anomalies in Colombia that exponentially increase dengue cases and extend dengue’s reach into high-altitude areas, potentially exposing millions of immunologically naïve people. This finding is highly relevant to the elevated dengue-free megacities in Latin America, such as Mexico City, Quito, and Bogotá. By shifting the focus from multi-decadal trends to interannual climate variability, this paper provides a methodological blueprint for more accurate predictions of how infectious diseases will behave as our climate becomes increasingly unstable.

## INTRODUCTION

Anthropogenic climate change presents an unprecedented challenge to humanity, with profound consequences for health and well-being (Romanello et al, 2025; IPCC, 2023). Among these impacts, changes in infectious disease dynamics stand out as a major concern, given evidence that climate change can expand the geographic and altitudinal range of climate-sensitive diseases and increase their epidemic potential (Alcayna et al 2025).

These climate-driven shifts in disease risk may be especially consequential for dengue in the Americas. Although dengue is endemic across the tropical lowlands of Central and South America, the risk distribution is strongly influenced by altitude. Brazil, with its densely populated urban centers at low to moderate elevations, such as São Paulo and Rio de Janeiro, experiences some of the highest dengue burdens worldwide (Junior et al., 2022). In contrast, countries like Ecuador, Peru, Colombia, and Mexico have historically been protected from major epidemics in their highest cities, such as Bogotá and Mexico City. Located above 2,000 meters, these urban centers are at elevations currently unsuitable for *Aedes* vectors. There is growing concern about the extent to which climate change will enable dengue to spread into previously protected high-altitude urban areas. Such outbreaks could be severe because of the large populations that are immunologically naïve and the demonstrated ability of *Aedes* mosquitoes to adapt to urban settings (Powell et al, 2013), potentially transforming the global landscape of dengue infections.

To date, the strongest empirical evidence linking climate change to altitudinal shifts in vector-borne disease comes from studies of highland malaria in Ethiopia and Colombia, which have demonstrated that the spatial distribution of cases shifts upslope during warmer years (Siraj et al., 2014). However, these foundational observations provide limited insight into the mechanisms that drive disease expansion into highland regions or the timescales over which such expansion occurs. This uncertainty is particularly important because climate change manifests through processes operating across multiple temporal scales, including changes in short-lived extreme events, interannual variability, and long-term warming trends (IPCC, 2023). Determining which of these processes most strongly influences the dengue elevational expansion remains an important unresolved question. In fact, current projections of disease expansion rely predominantly on mean-state trends, assuming that dengue will gradually encroach on high-altitude areas as global temperatures rise (Messina et al, 2019; Ryan et al, 2019; Colón-Gonzalez et al, 2021). However, before long-term projections can be formulated, it is essential to characterize the currently observed dynamics of the dengue altitudinal frontier and identify the primary drivers governing its fluctuations.

Colombia serves as an ideal testbed for this inquiry. It is a tropical country with endemic dengue transmission and a high concentration of densely populated urban centers distributed along the Andean elevational gradient. Currently, more than half of the Colombian population resides at elevations exceeding 1,000 m (IM Editores, n.d.). The country’s unique topography, where the Andean cordillera diverges into three distinct ridges, creates a natural experiment in which millions of people live at the suitability frontiers of transmission.

To date, Colombia exhibits a dual dengue transmission regime (Fig. S1). Lowland areas sustain endemic, year-round transmission governed by a complex interplay of climate, population immunity, sociodemographic factors, and vector control interventions. In the Andes, by contrast, transmission manifests as large but sporadic outbreaks that nevertheless account for more than half of the national burden. This epidemiological decoupling, whereby lowland and Andean dynamics differ substantially in their temporal structure, occurs at approximately 500 m elevation (Fig. S1). In highland regions, transmission operates near its lower thermal limit, population immunity remains low, and vector control is not a priority. Critically, unlike in the lowlands, where climate remains consistently within ranges permissive to transmission, even minor climatic anomalies can nonlinearly push these systems into conditions that trigger explosive outbreaks (Mordecai et al, 2017).

In this article, we analyze long-term epidemiological and meteorological data in the Colombian Andes to uncover the mechanisms and timescales that characterize dengue dynamics along an altitudinal gradient spanning from 500 m up to its elevational maximum at around 2000m. We demonstrate how interannual climate variability reorganizes both transmission intensity and the altitudinal range of epidemic risk in the Colombian Andes. To do so, we develop a mechanistic framework linking ENSO dynamics to local temperature anomalies, which in turn govern an upslope expansion of dengue into immunologically naïve highland populations. Beyond the Colombian Andes, this framework provides a general and reproducible approach for understanding disease dynamics near elevational range limits and for identifying the climatic drivers of altitudinal disease expansion.

Our findings challenge the paradigm that disease expansion is primarily a gradual, mean-state process. We show that interannual variability can be a dominant driver of both incidence and the geographic extent of infectious disease transmission. This has direct implications for how future risk should be projected, as frameworks that focus exclusively on mean-state trends risk mis-attributing or entirely missing dominant climate signals.

## RESULTS

### ENSO as the dominant driver of interannual climate variability

Previous studies have reported synchrony between ENSO and dengue dynamics in Colombia (Poveda et al., 2000; Poveda et al., 2001), suggesting that interannual climate variability plays a central role in shaping transmission. However, these studies remain largely observational, stopping short of mechanistically linking Pacific SSTs to their regional expression in Colombia and, further, to dengue transmission or altitudinal dynamics. Given these previous findings and the dominance of interannual variability in our dengue data, particularly in the 2–7 year ENSO band (Fig. S2), we first examined how ENSO structures interannual climate variability across the Colombian Andes.

We employed a hybrid data-driven approach combining Singular Spectrum Analysis (SSA) (Ghil et al, 2002; Golyandina et al 2018) and Empirical Orthogonal Function (EOF) analysis (see Materials and Methods). These methods facilitate the decomposition of raw monthly time series into distinct seasonal, interannual, and trend components and enable robust characterization of the dominant modes of spatiotemporal variability. We accessed reanalysis data (see Materials and Methods) and applied SSA as a temporal filtering technique to isolate the interannual component of temperature and precipitation at each grid point. Then, EOF analysis was performed on the SSA-reconstructed interannual fields to extract the dominant spatial patterns and their associated principal components. This workflow enables characterization of the spatiotemporal variability of the interannual component of temperature and precipitation.

This sequential approach is important because an EOF run on the raw signal would maximize total variance across all timescales, thereby conflating variability associated with distinct physical mechanisms. By applying SSA beforehand, we empirically separate components associated with different timescales, allowing the EOF decomposition to be performed specifically on the interannual signal rather than on the full mixed-timescale field. This ensures that the resulting spatial modes primarily reflect interannual variability most relevant to dengue dynamics.

This analysis revealed that the climate in the Colombian Andes is strongly influenced by ENSO (Fig. 1A, 1B). This is confirmed by the close correspondence between the interannual temperature and rainfall PCs, which track the Oceanic Niño Index (ONI) signal (Fig S1B, S2). Both PCs show a very high Spearman’s rank correlation (rho > 0.85) with each other when the temperature PC is lagged by two months relative to the rainfall PC (Fig. S3). This indicates that the effect of ENSO first manifests in rainfall (causing drought during the warm phase), with the peak warming effect occurring two months later. This lag is consistent with the Walker Circulation’s faster response relative to the slower, ocean-driven buildup of regional temperatures.

**Figure 1.**
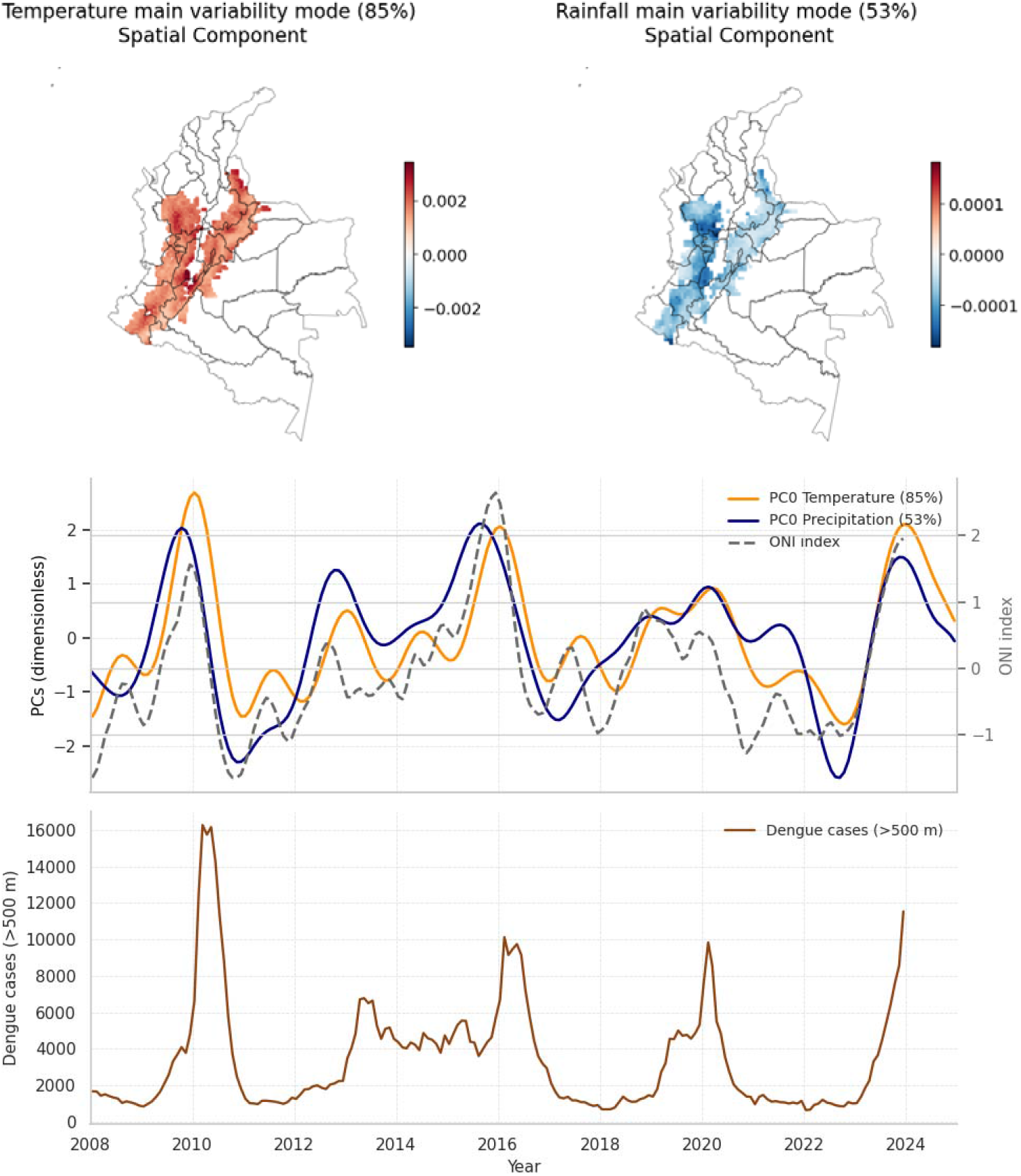
ENSO influence on interannual climate variability in the Colombian Andes and its relationship with dengue transmission. **(A)** Spatial patterns (empirical orthogonal functions, EOFs) of the leading modes of interannual variability in temperature and precipitation over the Colombian Andes, derived from gridded ERA5 (temperature) and CHIRPS (rainfall) data after decomposition using Singular Spectrum Analysis (SSA). **(B)** Corresponding temporal components (principal components, PCs) associated with the leading EOFs of temperature and precipitation, shown together with the Oceanic Niño Index (ONI). Each mode of variability is represented by a combination of a spatial pattern (EOF) and a temporal component (PC); the reconstructed field is obtained by multiplying the EOF by its corresponding PC. In this case, a similar PC with oppositely signed spatial patterns—positive anomalies for temperature (red) and negative anomalies for precipitation (blue)—indicates contrasting climatic responses during ENSO phases. **(C)** Monthly dengue case counts reported at elevations above 500m across Colombia, shown for comparison with the PCs and ONI time series in panel B.

With this workflow, we extract the ENSO signal from both local temperature and rainfall data in Colombia, accounting for ∼85% and ∼53% of total interannual variability in temperature and rainfall, respectively. The extracted PC, along with its associated spatial patterns, corresponds to the regional ENSO manifestation in Colombia, or in more climatic terms, the ENSO local teleconnection.

Dengue incidence closely mirrors the PCs/ONI signal (Fig. 1B vs. 1C or Fig. S4), suggesting it is strongly ENSO-modulated. Physically, ENSO drives both generalized warming and drought across the Colombian Andes. Elevated temperatures are a well-understood driver of dengue transmission, as they accelerate the disease transmission cycle. Conversely, while drought might intuitively be expected to limit mosquito populations by desiccating natural breeding sites, empirical evidence in Colombia and similar tropical contexts suggests it can also favour them if water storage practices are widely adopted during droughts (Lowe et al., 2021; Kache et al., 2024). This behavioral adaptation could offset the biological constraints of drought, so it is necessary to formally disentangle whether dengue dynamics are primarily driven by ENSO-driven increases in temperature or by reduced rainfall.

#### Synchrony between ENSO and dengue altitudinal expansion

After observing the strong link between dengue incidence and ENSO-driven local climate variability, we further investigated whether ENSO also modifies the altitudinal distribution of transmission. A preliminary analysis comparing the altitudinal distribution of dengue incidence during El Niño months with that during La Niña months (defined according to NOAA ENSO criteria as periods with an ONI index > 0.5 (or < −0.5 for La Niña) lasting at least five consecutive months) revealed significant differences in the distribution of cases between these ENSO phases (Fig. 2A).

**Figure 2.**
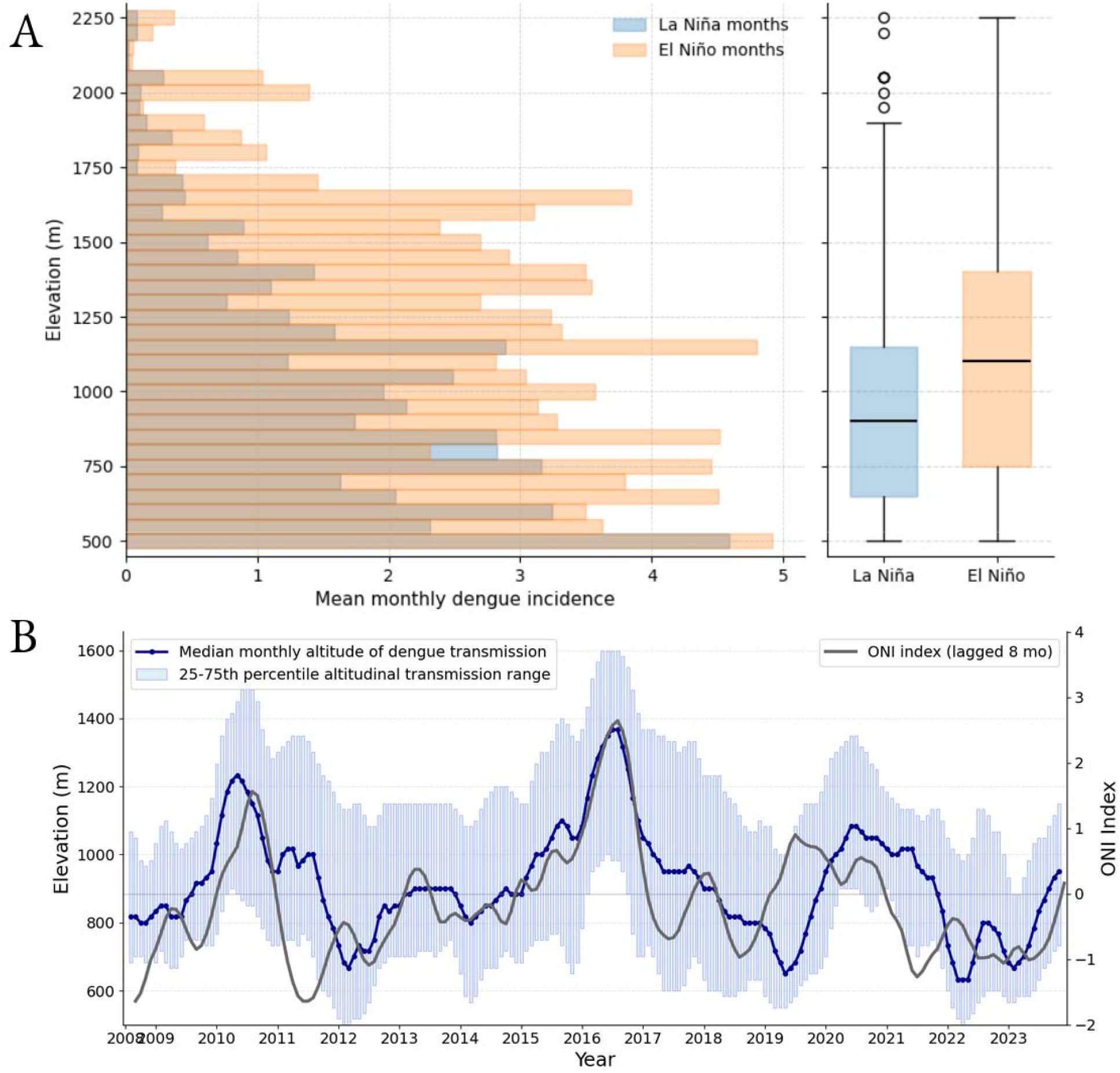
Altitudinal distribution of dengue incidence and its temporal dynamics in the Colombian Andes. **(A) Left:** Histograms showing the mean monthly dengue incidence as a function of elevation during different ENSO phases, illustrating both higher incidence levels and an upward shift in the altitudinal distribution during El Niño months compared with La Niña months. **Right:** Boxplots summarizing differences between the elevation distribution of cases during El Niño months and La Niña months. El Niño and La Niña months were identified according to NOAA ENSO criteria, defined as periods with an ONI index > 0.5 (El Niño) or < −0.5 (La Niña) that lasted at least 5 consecutive months. An 8-month lag was applied to the ONI index, corresponding to the maximum cross-correlation delay identified between ONI and the altitudinal distribution of dengue cases. This lag reflects a sequential cascade: Pacific SST anomalies propagate to local Colombian temperature with a delay of approximately 2–3 months (Fig. S2), which in turn influences dengue transmission and the upslope expansion of cases over the following months. See the “Lag structure of the mechanistic cascade” section for a thorough derivation of the component lags underlying this estimate. **(B)** Temporal evolution, at monthly resolution, of the median elevation (Q2) at which dengue cases are reported, along with the interquartile range (Q1, Q3) of the dengue altitudinal distribution (as in panel A, right), shown together with the Oceanic Niño Index (ONI) lagged by 8 months.

Specifically, during La Niña months, the first quartile of the altitudinal distribution of dengue cases is at approximately 680 m, and the third quartile at approximately 1,150 m. During El Niño (warm) phases, the lower quartile shifts upward to approximately 750m and the upper quartile to 1,400 m, indicating an upward displacement of the dengue transmission zone (Fig. 2A).

To further investigate this dynamic, we examined the monthly evolution of the distribution’s quartiles. This analysis revealed a coherent temporal progression across all quartiles that closely tracks the ONI signal (Fig. 2B).

### ENSO spatial signatures and connection to dengue

Having identified a strong synchrony between the interannual PCs of temperature and precipitation and both dengue incidence and its altitudinal distribution, we next sought to disentangle the respective roles of these variables and identify the primary driver. However, high collinearity between the temperature and precipitation PCs limits their robust separation in the temporal domain alone. We therefore first examined the spatial signatures of ENSO teleconnections.

In Colombia, the Andes consist of three ridges (Western, Central, and Eastern), separated by the Cauca and Magdalena river valleys. For simplicity and in line with standard practice, we classified municipalities into two main categories: Western and Eastern ridges, using the Magdalena River Valley as the dividing boundary (Fig. 3A). Our SSA-EOF analysis revealed an informative spatial divergence: while both ridges exhibit uniform sensitivity to ENSO-driven temperature fluctuations (Fig. 3B), the Western ridge shows significantly higher sensitivity to ENSO-driven rainfall oscillations (Fig. 3C). This is supported by pixel-level distribution analysis of the rainfall EOFs across the two ridges, which reveals distinct distributions that are statistically significant under a bootstrap randomization test.

**Fig 3.**
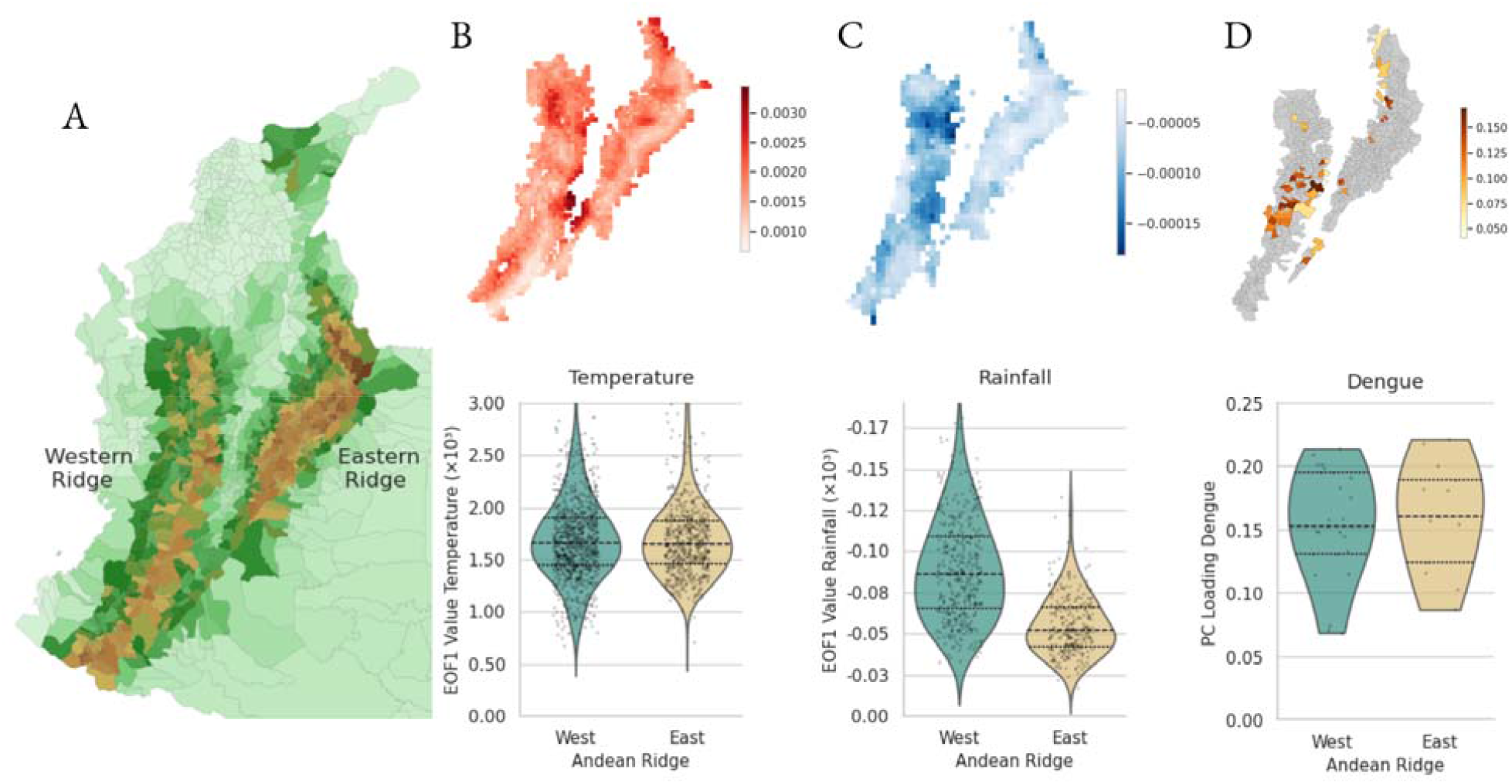
ENSO and dengue spatial signatures. (A) Geographic profile of Colombia, highlighting the division of the Andes into two distinct mountain ridges separated by the Magdalena Valley. (B) Temperature EOF patterns across the ridges (top) and distribution of EOF values by ridge (bottom), shown as full scatter plots (black dots), violin plots, and summary statistics including the median (dashed line) and Q1 and Q3 quartiles (solid lines). (C) Same as in (B), but for rainfall, showing clear differences between ridges. (D) Same as in (B), but for dengue incidence, including only municipalities reporting more than five cases per month to ensure robustness and minimize stochastic noise. Sensitivity analyses using thresholds ranging from 1 to 10 minimum monthly cases are presented in Fig. S5.

These findings are consistent with known ENSO dynamics from a physical perspective. The Western ridge, located near the Pacific coast, experiences more direct modulation of moisture transport and is more sensitive to changes in the Walker circulation. Meanwhile, the Eastern ridge, farther from the ocean and protected by the Western ridge, shows reduced ENSO influence on rainfall. In contrast, temperature changes are driven by a slower, more uniform regional warming, leading to the more consistent spatial pattern seen in our results.

To study the regional patterns of dengue, we performed a Principal Component Analysis (PCA) at the municipal level, stratified by ridge. Given the inherent stochasticity of municipal-level data and PCA’s sensitivity to noise, we standardized the incidence rates and employed an iterative PCA approach. By applying a selection threshold (ranging from 1 to 10 minimum monthly cases), we filtered out low-incidence municipalities to prevent regions with a low signal-to-noise ratio from obscuring or diluting the underlying ENSO signal. Across all tested thresholds (see Supplementary Materials), dengue incidence exhibited similar ENSO sensitivity across both ridges (Fig 3D). This spatial homogeneity closely aligns with the observed temperature patterns but diverges from the ridge-specific rainfall pattern.

### Nonlinear temperature effects on dengue dynamics

We then moved to the time domain to reassess which variables (local or global) better account for dengue incidence and elevation dynamics. To do so, we compared several Generalized Additive Model (GAM) formulations for both dengue incidence and the elevation at which cases are reported. Given the high collinearity among interannual temperature and precipitation components and the Oceanic Niño Index (ONI) (Fig. 1B), we could not simultaneously incorporate these variables. We therefore employed univariate models alongside bivariate models using orthogonalized residuals (e.g., local temperature plus the component of the ONI signal not already captured by local temperature). Model selection was performed by balancing in-sample fit (AIC, Pseudo R^2^), out-of-sample validation (MAE, CV Pseudo R^2^), and model stability (Δ Pseudo R^2^), as detailed in Table 1. Out-of-sample validation was done using a forward-chaining cross-validation approach.

**Table 1.** Comparison of dengue incidence (A) and dengue elevation (B) model performance and stability. Models are different Generalized Additive Model (GAM) formulations, specified in the Model Structure column, and fitted using a Poisson likelihood with a log link for case counts. Smooth terms were constrained with 2 knots of degree 2. Variables with the suffix *_res* denote orthogonal residuals (e.g., rainfall signals independent of temperature), used to reduce collinearity.

| Model Structure<br>(Predictor Variables) | In-Sample (Full Fit) |  | Out-of-Sample (CV) |  | Stability |
| --- | --- | --- | --- | --- | --- |
| | Ps. $R^2$ | AIC | Ps. $R^2$ | MAE | Drop ($\Delta R^2$) |
| temp | 0.76 | 87041.66 | **0.66** | **762.13** | **0.10** |
| temp + oni_res | 0.76 | 85233.12 | 0.64 | 868.75 | 0.12 |
| temp + rain_res | **0.79** | **74806.56** | 0.52 | 936.59 | 0.27 |
| rainfall + temp_res | **0.79** | 76475.88 | 0.51 | 968.74 | 0.28 |
| rainfall + oni_res | 0.73 | 98176.83 | 0.49 | 984.52 | 0.24 |
| rainfall | 0.69 | 111268.99 | 0.42 | 968.12 | 0.27 |
| oni | 0.64 | 130534.76 | 0.42 | 1156.54 | 0.22 |

**Table 1: Comparison of dengue incidence (A) and dengue elevation (B) model performance and stability.**
| Model Structure<br>(Predictor Variables) | In-Sample (Full Fit) |  | Out-of-Sample (CV) |  | Stability |
| --- | --- | --- | --- | --- | --- |
| | Ps. $R^2$ | AIC | Ps. $R^2$ | MAE | Drop ($\Delta R^2$) |
| temp | 0.51 | 2323.69 | **0.44** | **98.28** | **0.07** |
| rainfall + temp_res | **0.56** | 2104.01 | **0.44** | 102.08 | 0.14 |
| rainfall | 0.52 | 2266.61 | 0.42 | 108.30 | 0.10 |
| temp + rain_res | 0.52 | 2272.69 | 0.38 | 106.65 | 0.14 |
| temp + oni_res | **0.56** | **2083.35** | 0.36 | 106.57 | 0.20 |
| oni | 0.45 | 2606.30 | 0.28 | 116.21 | 0.17 |
| rainfall + oni_res | 0.53 | 2231.33 | 0.22 | 124.62 | 0.31 |

In both models (Table 1A), a univariate temperature model outperforms all other formulations in predictive skill, achieving the lowest MAE, the highest CV Pseudo R2, and the lowest R2 drop between fits and CV. Figure 4 illustrates the fits of the best-performing models and the corresponding partial effects of temperature. The response functions reveal pronounced nonlinearities, with both dengue incidence and altitudinal limits increasing exponentially at high PC values of temperature, corresponding to ENSO warm phases.

**Fig. 4.**
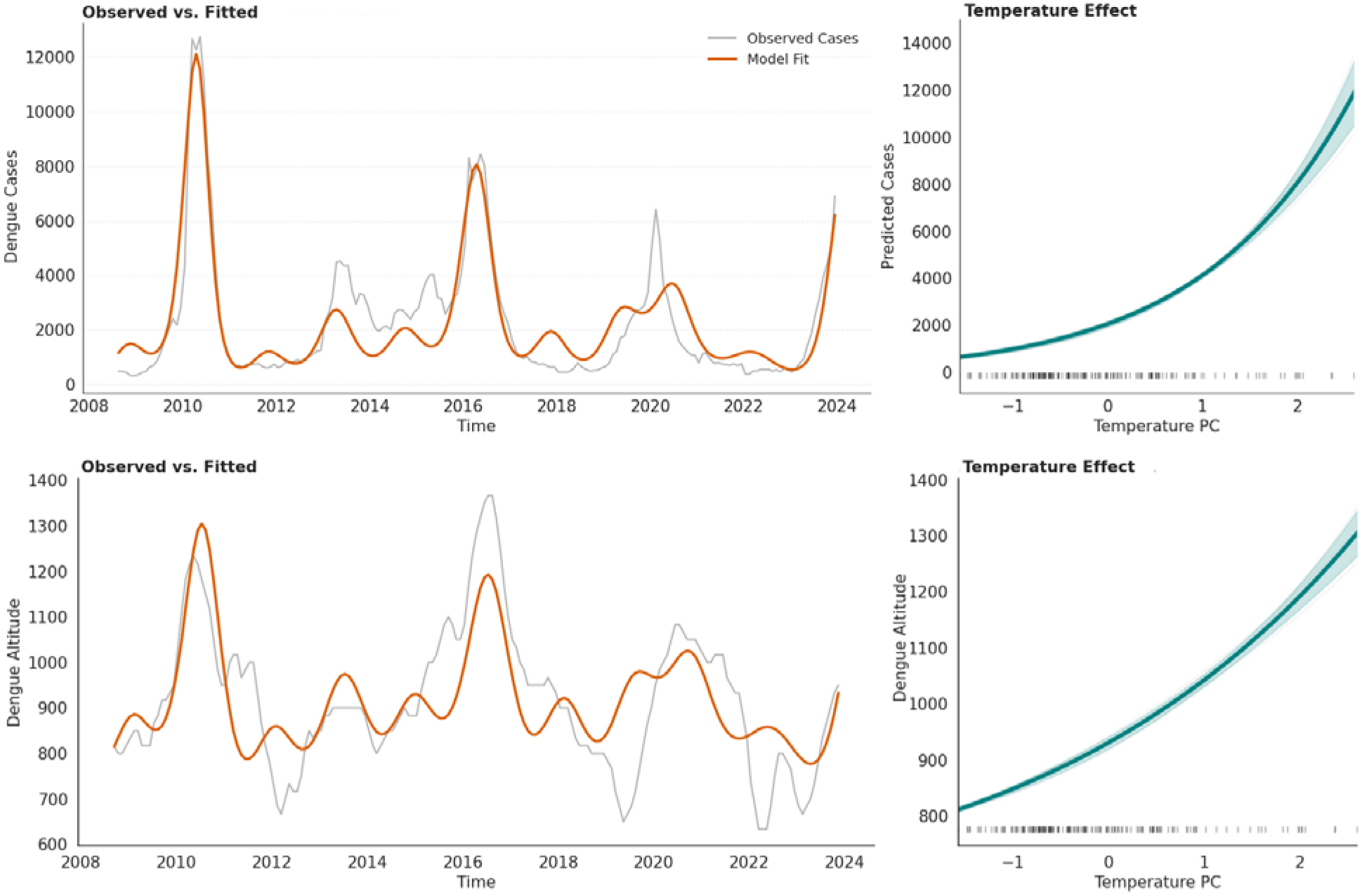
Top-performing model fits and response functions. Panel (A) shows the univariate temperature-driven incidence model, with its corresponding dose-response function shown on the right. Panel (B) shows the univariate elevation model driven by temperature, with its corresponding dose-response function displayed on the right.

Importantly, the results also confirm that once local climate is properly accounted for, ENSO indices provide no additional explanatory power. While the inclusion of ONI marginally improved in-sample metrics, it consistently degraded out-of-sample performance and reduced model stability. This supports the “mechanistic cascade” hypothesis: ENSO modulates local interannual temperature and precipitation, which, in turn, directly drives dengue. While ENSO indices and local principal components (PCs) share significant variance, local indices likely capture specific environmental nuances that global indices do not, making them superior predictors of regional transmission.

#### Mechanism of altitudinal expansion

Dengue incidence and its altitudinal expansion dynamics are strongly synchronized, both increasing during warm ENSO phases and decreasing during cold ones. They also show similar exponential-like temperature dose-response functions (Fig. 4). This synchrony raises a fundamental question: does altitudinal expansion emerge from overall increases in incidence, or does ENSO generate elevation-specific effects that independently shift the transmission zone upslope?

ENSO-related temperature anomalies are relatively coherent across the altitudinal gradient of our study region (Yulaeva et al, 2004; also Fig. S6), suggesting that elevation-specific temperature effects are unlikely to be the primary driver of such coupling. To investigate how increases in incidence could translate into altitudinal expansion, we employed a temperature-dependent mechanistic model of mosquito life-history traits to estimate relative vectorial capacity (rVC). We used the standard formulation (Kramer et al, 2015; Mordecai et al, 2019):

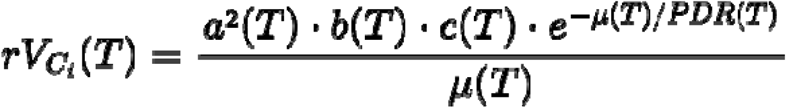

where a(T), b(T), c(T), PDR(T) and \mu(T) denote the biting rate, the transmission probability, the infection probability, the parasite development rate, and the mortality rate, respectively; each modeled as a function of temperature using thermal response functions from Mordecai et al. (2019), parameterized for *Aedes aegypti* and dengue transmission (see Appendix for further details). Intuitively, relative vectorial capacity can be interpreted as the temperature-dependent transmission efficiency of the vector population.

Using daily observed temperature time series, we estimated vectorial capacity for several Colombian Andean cities *i* along an elevational gradient: Bucaramanga (∼950 m), Cali (∼1000 m), Pereira (∼1410 m), Medellín (∼1500 m), Armenia (∼1550 m), Popayán (∼1740 m), and Bogotá (∼2600 m). These major urban centers cover the two main ridges and offer a representative cross-section of the altitude gradient. The goal is to illustrate mechanisms across different thermal environments rather than to provide exhaustive spatial coverage.

We computed the elevational centroid of vectorial capacity as:

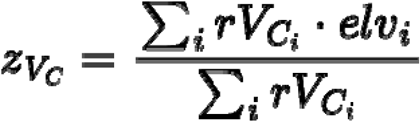

and further standardized it (using z-scores) to focus strictly on its dynamics. This index tracks the temporal evolution of the mean elevation at which dengue temperature-driven transmission potential is concentrated.

Strikingly, isolating this single component of the transmission process reveals that vectorial capacity alone reproduces the temporal dynamics of the median elevation at which observed dengue cases are reported (Fig. 5B). The mechanistic model accounts for >50% of the variability in observed altitudinal shifts (Spearman r^2^=0.52, Pearson r^2^=0.55, both p<0.01), without any parameters fitted to the altitudinal data, and setting aside human susceptibility, population distribution, and socioeconomic factors.

**Fig 5.**
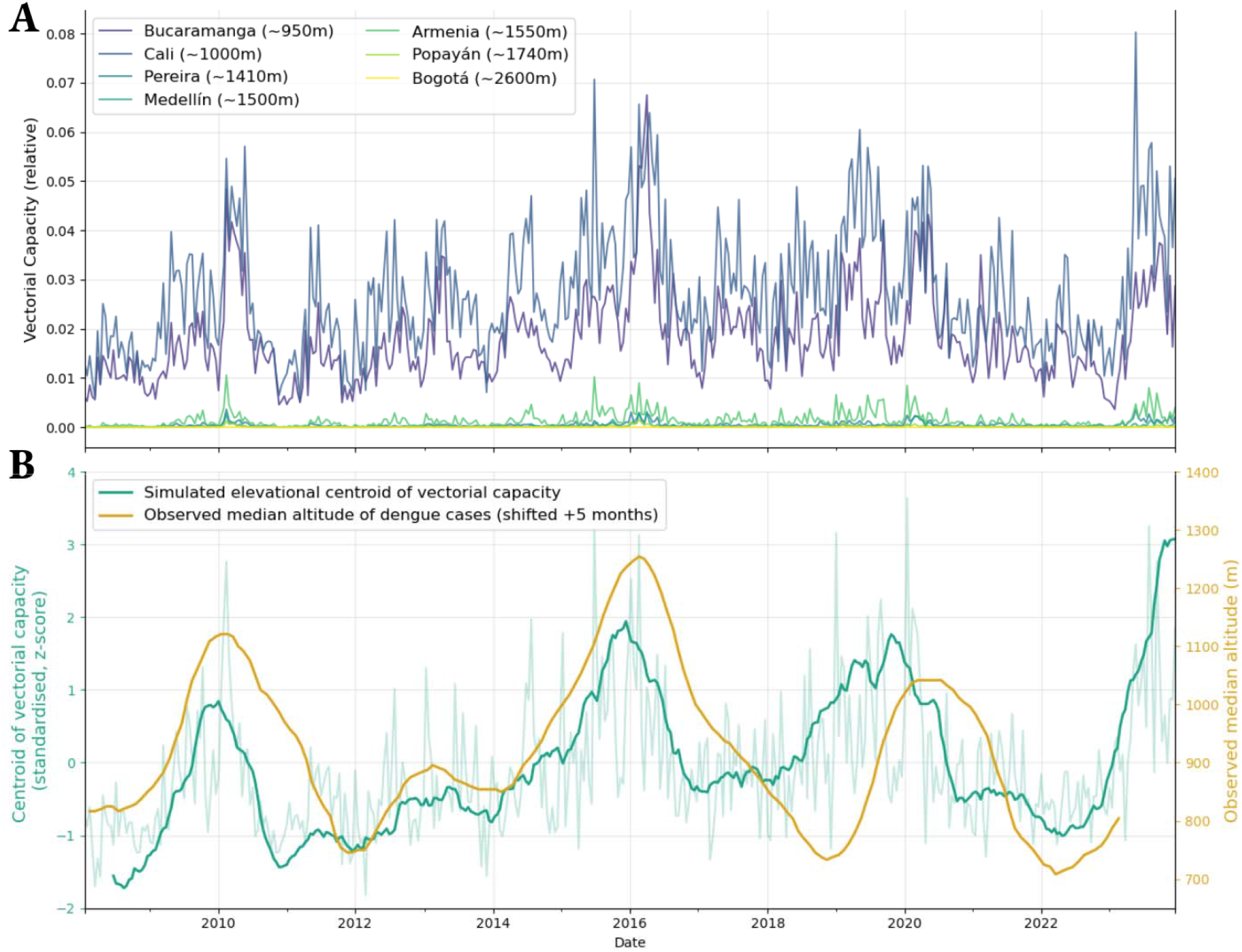
Mechanism of altitudinal expansion. (A) Temperature-driven simulations of vectorial capacity across seven representative Colombian Andean cities spanning an elevational gradient. Observed daily temperature time series are used as input to a mechanistic model of mosquito transmission potential, resampled to bimonthly frequency to improve visual inspection. (B) Simulated elevational centroid of vectorial capacity compared against the observed median elevation of reported dengue cases, both smoothed with a 12-month running mean to remove seasonality and focus on interannual dynamics. The dimmed lines show the underlying biweekly signal prior to smoothing.

This result implies that elevational shifts in dengue transmission can be explained, in large part, by temperature-driven changes in the distribution of vectorial capacity, without the need of invoking other covariates.

While lower-elevation regions often fall near the thermal optimum for transmission, higher-elevation fringe regions lie on the steeper, more sensitive portion of the thermal response curve. Consequently, a relatively uniform increase in temperature across elevational strata disproportionately enhances transmission potential in these previously marginal high-altitude zones, effectively lifting the transmission centroid upward.

#### Lag structure of the mechanistic cascade

The cascade from Pacific SSTs to dengue spread in highland areas unfolds through a series of sequential lags. Such lags, however, are variable and scale with forcing strength: for example, strong ENSO phases drive rapid, coherent responses across the cascade, while weak phases produce longer or undetectable effects; consistent with ENSO acting as an intermittent rather than continuous forcing. Having identified the mechanisms operating at each transition, the mean peak coupling lags can now be identified by cross-correlation analysis. ONI leads local temperatures in Colombia by approximately 2 months, and local temperature leads dengue incidence by approximately 3 months (Fig. S7). These delays are consistent with the time required for Pacific SST anomalies to drive atmospheric warming in Colombia, and for that warming to accelerate mosquito life-history traits, shorten the extrinsic incubation period, and produce detectable increases in reported cases. However, the effect of temperature on incidence is not uniform across elevational strata. Colder high-altitude zones experience a delayed onset of outbreaks relative to lower elevations, as the same warming anomaly takes longer to cross transmission thresholds there. This is illustrated by Fig. S8A, which compares incidence dynamics between Cali (∼1,000 m) and Medellín (∼1,550 m), showing that epidemic peaks in the higher-altitude city consistently lag those at lower elevation. Fig. S8B provides a closer look at the 2010 ENSO-driven outbreak revealing a progressive delay in transmission buildup with increasing altitude.

The median elevation at which dengue cases are reported is affected by these differential outbreak timing across the altitudinal gradient and presents larger lags with temperature than incidence, which is a metric dominated by lower elevations where most cases occur. Consistent with this, cross-correlation analysis identifies a peak coupling between local temperature and the median elevation of dengue cases at approximately 5–6 months, depending on whether the mechanistic vectorial capacity model or the phenomenological GAM is used as reference. Combined with the ∼2-month ONI-to-temperature lag, this yields a cumulative delay between ONI and elevational patterns of approximately 8 months, broadly consistent with the sum of these component delays and supporting a sequential causal pathway from Pacific sea surface temperatures through local climate to transmission intensity and geographic expansion.

This sequential timing defines a predictable propagation pathway from Pacific climate forcing to local epidemic dynamics, one that can be operationalised for early warning systems and that provides a principled basis for attributing disease risk to large-scale climate drivers.

#### Convergence of evidence

We provided converging evidence from multiple independent analyses supporting a mechanistic cascade linking ENSO variability to dengue transmission through local temperature. First, we showed that interannual climate variability in the Colombian Andes is strongly ENSO-modulated, synchronizing both dengue incidence and its altitudinal expansion. To disentangle the respective roles of temperature and precipitation, we examined the spatial structure of ENSO teleconnections and dengue variability. Temperature-related anomalies exhibited coherent spatial patterns closely aligned with dengue dynamics, whereas precipitation displayed more heterogeneous, ridge-specific responses that were not mirrored in dengue incidence patterns.

Second, time-domain analyses consistently identified temperature as the dominant predictor of both dengue incidence and altitudinal shifts. Across all candidate formulations, precipitation did not improve out-of-sample predictive performance once temperature was accounted for and was not retained under cross-validation model selection criteria.

Third, the mechanistic vectorial capacity analysis reproduced a substantial fraction of the observed altitudinal variability using temperature-dependent mosquito traits alone through non-linear responses. With this, two key empirical features of the system can naturally be reproduced: the strong synchrony between incidence and altitudinal expansion, and also the delayed epidemic response observed in colder high-altitude regions relative to warmer lower-elevation areas. Both patterns are results of nonlinear thermal constraints on transmission and are difficult to reconcile with precipitation forcing.

These results suggest that precipitation contributes very limited independent explanatory power for interannual dengue variability in the Colombian Andes at the spatial and temporal scales considered here. In contrast, the results consistently support temperature as the dominant proximate driver of both transmission intensity and upslope expansion. Given that these highland regions lie near the lower thermal limit for dengue transmission, such strong temperature sensitivity is biologically and climatologically consistent.

## DISCUSSION

### Implications for climate-disease scenario analysis

The mechanistic cascade demonstrated here, from ENSO to altitudinal expansion, has direct consequences for how past and future climate-sensitive infectious diseases scenarios are generated. While anthropogenic forcing alters both the mean climate and modes of variability, substantial uncertainty remains about how these modes will change under future warming, particularly for coupled ocean–atmosphere phenomena such as ENSO and the NAO (Liu et al., 2025; Mitevsky et al., 2025). As a result, many current detection and attribution (D&A) approaches in infectious diseases focus on long-term trends or extreme events, while implicitly downplaying seasonal and interannual variability (Carlson et al., 2024). This limits their applicability in systems where variability is the primary driver of transmission dynamics. Our results suggest that explicitly incorporating climate variability is essential for robust attribution studies of climate-sensitive infectious diseases, as omitting it risks discarding the most relevant signals.

A key contribution of this study is that we ground dengue dynamics in the local expression of ENSO teleconnections rather than in remote sea surface temperatures (SSTs). This shift has important implications for counterfactual scenarios and future projections. While significant uncertainty remains regarding how ENSO frequency and amplitude will respond to anthropogenic forcing (CMIP6 models diverge on whether Pacific variability will increase or decrease), there is greater model consensus that some teleconnections may intensify (Trascasa-Castro et al., 2025). This is because, in a warmer world, thermodynamic amplification may enhance hydrological responses to ENSO-related circulation anomalies. Therefore, in the case of Colombia, projections and detection and attribution analyses may be more feasible when grounded in local teleconnection responses, which may exhibit greater consistency across models while ENSO’s future behaviour remains uncertain. This is a possible way forward that will be explored in subsequent studies.

Finally, our findings challenge a common assumption in current disease range projection frameworks, which typically rely on changes in mean climate conditions under anthropogenic warming. We show instead that the spatial footprint of dengue in the Colombian Andes is highly sensitive to interannual variability, with climate oscillations capable of rapidly reorganizing the altitudinal distribution of transmission on much shorter timescales than those associated with long-term trends. This suggests that approaches based solely on mean-state changes may underestimate the role of climate in driving disease risk. Explicitly incorporating climate variability into predictive and attribution frameworks is therefore essential for accurately anticipating future epidemic risk in a warming world.

#### Implications for public health

The sequential cascade identified here, linking Pacific SST anomalies to local temperature to transmission intensity, and geographic expansion, unfolds over a predictable ∼8-month window. ENSO forecasts are routinely issued several months in advance with meaningful skill, so the full ONI-to-epidemic delay is potentially forecastable end-to-end, providing ample time to deploy targeted interventions. This creates a rare and valuable opportunity: the main driver is global, well monitored and predictable; the pathway to epidemics is now better understood; and the populations most at risk can be identified and targeted geographically.

This stands in contrast to lowland dengue transmission in Colombia, where endemic year-round dynamics are governed by a complex interplay of population immunity, serotype circulation, sociodemographic factors, and vector control, making forecasting substantially harder. Highland Andean transmission, by contrast, is episodic and strongly climate-driven, with interannual variability accounting for the dominant share of epidemic risk. This simplicity is a major asset: the signal is strong, the noise is relatively low, the lead time is long..

A developing El Niño should therefore serve as an operational trigger for Andean cities: a coordinated protocol combining intensified entomological surveillance, preemptive vector control, and public health communication could substantially reduce epidemic impact at a fraction of the cost of reactive responses.

### Local climate teleconnections outperform global ENSO indices

A significant body of literature on infectious disease dynamics (Hallett et al, 2004; Chaves et al, 2006) relies on analyses of raw climate data alongside ENSO indices, often concluding that the latter have superior predictive power. This has fostered a widespread consensus that ENSO indices may capture the “true” climate signal more effectively than local station or gridded data (Hallet et al, 2004; Stenseth et al, 2002). Our results question the usual assumptions, suggesting that these conclusions may not accurately reflect the underlying climate dynamics. Instead, they could be artifacts caused by inadequate time-series decomposition methods.

This study employed a combination of Singular Spectrum Analysis (SSA) and Empirical Orthogonal Functions (EOFs), a suite of nonparametric, data-driven techniques specifically designed to handle the nonstationary and anharmonic nature of climate signals. They allowed us to extract the local ENSO teleconnections and ground dengue dynamics in the local expression of ENSO rather than in Pacific SSTs. This strategy provides a clearer mechanistic sequence, linking ultimate drivers like ENSO to proximate factors such as local climate, and then to the transmission process. Since disease spread is a localized event, the fact that Pacific Sea Surface Temperatures (SSTs), thousands of kilometers away, often predict outcomes better than local conditions presents a biological paradox that deserves further investigation (Stenseth et al, 2003).

With the exception of the Southern Oscillation Index (SOI), most widely used ENSO indices (e.g., ONI, Niño 1.2, etc) are oceanic indices derived from Pacific SSTs. Due to its high inertia, the ocean acts as a natural low-pass filter for climatic variability. In contrast, local weather, arising from a blend of processes operating at subseasonal, seasonal, and interannual scales, merges multiple sources of variability that blur the ENSO signal. In practical terms, this typically implies that the ENSO teleconnection is buried within strong seasonal climatic variability, which may explain why oceanic indices have historically outperformed raw local data in predictive models.

In addition to providing a more biologically sound explanation of how ENSO forcing operates, the approach outlined here has several advantages, including greater mechanistic clarity and improved predictive optimization. Ultimately, prioritizing the extraction of local signals over broad global indices provides both theoretical rigor and practical relevance for public health surveillance.

#### Limitations

The main limitation of our study is that it relies on reported dengue cases for which the place of infection is assumed to be correctly identified. Misclassification may arise from reporting errors, travel-associated infections, or uncertainty about the location of exposure. Also, municipalities with multiple population centers at different elevations could bias our altitudinal attribution. Despite this limitation, this study analyses the full elevation distribution and identifies shifts across all quartiles. These metrics are stable and minimally sensitive to sporadic misclassification, which is expected to affect only a small fraction of cases. We deliberately refrained from analyzing maximum elevation, a metric particularly sensitive to outliers and reporting errors.

Another limitation of this study is the lack of data on human mobility between lowland and highland cities, which prevents us from directly testing the hypothesis that dengue epidemics originate in endemic lowland regions and are later introduced into highland cities by infected travelers, leading to delayed epidemic onset at higher elevations. Such importation events likely help seed transmission in highland locations. However, they may not by themselves explain the gradual upslope progression of epidemic peaks observed, for example, during the 2010 outbreak. A purely mobility-driven mechanism would not necessarily produce systematic delays in peak timing with increasing elevation. Instead, the observed pattern is more consistent with a gradual increase in suitable transmission conditions as temperature becomes more favorable along the elevational gradient.

Importation pressure from lowland to highland regions is expected to scale with dengue incidence in lowland areas and therefore vary substantially over time. However, introductions alone cannot determine whether outbreaks amplify once cases arrive in highland settings. Instead, local transmission appears to be primarily constrained by local climatic suitability, particularly temperature-dependent limits on vectorial capacity. In this framework, human mobility determines seeding events, while temperature governs amplification. The decoupling between lowland endemic dynamics (so likely also importation pressure) and highland epidemic timing therefore suggests that climate is instead the dominant determinant of highland dynamics, although the relative contribution of seeding and local environmental constraints cannot be properly quantified without mobility data.

Empirical studies indicate that *Aedes aegypti* abundance and dengue transmission decline sharply above ∼2,000 m (Lozano-Fuentes et al, 2012). Sporadic vector detections have been reported at higher elevations, including up to ∼2,550 m in Bolivia (Castillo-Quino et al, 2018; Equihua et al, 2017), but these are rare and have not been associated with dengue transmission. In Colombia, the highest reported *Aedes* mosquito was observed at ∼2,300 m, and dengue-infected *Aedes* at ∼2,000 m (Ruiz-López et al., 2016). In our dataset, only 0.17% of cases are reported above 2,300 m. Given their rarity and the increased likelihood of reporting error, we excluded these cases from Fig. 2a for visualization purposes. Fig. S9 reproduces Fig. 2 using the complete dataset, confirming that this elevation threshold does not affect our conclusions. Despite these caveats, the convergence of spatial, temporal, and mechanistic evidence presented in the Results makes the principal findings robust to these uncertainties.

## CONCLUSIONS

This study shows that the altitudinal boundary of dengue is not a fixed ecological limit but a dynamic climate-structured frontier that expands during warm ENSO phases. Through a mechanistic cascade linking ENSO to local temperature variability and, via nonlinear transmission responses, to incidence and altitudinal range, we demonstrate that interannual climate variability rapidly reorganizes the geography of epidemic risk. These dynamics operate on timescales far shorter than those associated with long-term warming trends.

Rather than modulating incidence within a stable spatial domain, ENSO-driven variability shifts both the intensity and upper bound of transmission, periodically exposing high-altitude populations that are otherwise below the epidemic threshold. This creates a predictable window of elevated risk that can be operationalized: ENSO forecasts could provide actionable lead times for preemptive vector control and resource allocation in high-elevation cities.

Unlike most studies of geographic disease expansion, which rely on vector suitability indices to project future potential ranges, this study is grounded in confirmed case data and documents observed, present-day altitudinal shifts. That these shifts are largely explained by interannual temperature variability, rather than long-term trends, fills a critical empirical gap in the literature and offers a methodological blueprint that can be extended to other climate-sensitive diseases and regions.

The highly urbanized Colombian Andes provided an ideal setting for using high-resolution monthly municipal data to examine the mechanisms and timescales of altitudinal expansion. However, the underlying mechanism is likely generalizable to other high-elevation tropical regions influenced by ENSO or analogous modes of climate variability.

Finally, and more broadly, this work challenges current scenario analyses, including Detection and Attribution studies in infectious disease systems. Robust risk assessment requires moving beyond multi-decade mean-state climate trends to explicitly incorporate interannual climate variability and its teleconnections. In a warming world, changing climatic variability as well as long-term warning trends will define the emergence and expansion of infectious diseases.

## MATERIALS AND METHODS

### Data and Elevation Determination

Monthly dengue data from 2008 onward were obtained at the municipal level in Colombia (1,122 municipalities) and served as the basis for studying incidence and elevation dynamics. Since dengue is an urban disease transmitted in population centers, we calculated, for each municipality, a population-weighted elevation to accurately capture the elevation where people actually live. We used WorldPop data and a digital elevation model (DEM) from the Shuttle Radar Topography Mission (SRTM) for this calculation. We further focused on Andean transmission (>500m) to avoid endemic areas where immunity can buffer or amplify the climate signal and to achieve a more climate-driven system while still representing half of the national burden.

Temperature data were obtained from the ERA5 reanalysis product via the Copernicus Climate Data Store. For precipitation, a more stochastic and spatially heterogeneous variable (thus showing higher observational uncertainty), we evaluated two widely used gridded datasets: ERA5 and CHIRPS. Both products were validated against in situ station data across Colombia (Fig. S10). Our comparative analysis indicated that, while both datasets adequately capture low-frequency variability in the Andean region (the primary focus of this study), CHIRPS shows better agreement with station observations (Fig S11). We therefore used CHIRPS precipitation data in all subsequent analyses.

### Singular Spectrum Analysis (SSA)

Singular Spectrum Analysis (SSA) is a data-driven, non-parametric method for decomposing a time series (e.g., disease incidence, climate variables) into orthogonal components. These components can then be grouped and reconstructed into meaningful signals such as trends, oscillatory modes, or noise.

The method works in three main steps:

1. **Embedding:** The original time series is mapped to a trajectory matrix by sliding a fixed-length window along it.
2. **Decomposition:** Singular value decomposition (SVD) is applied to this matrix, producing orthogonal elementary components.
3. **Grouping & Reconstruction:** Elementary components are combined into interpretable signals (e.g., seasonal cycles, interannual variability, long-term trends).

SSA is particularly useful for identifying oscillatory patterns and disentangling overlapping physical or biological processes, even in short or noisy datasets. We used the RSSA package, which includes a periodogram-based grouping algorithm to help classify components into interpretable signal types. Grouping is semi-automated but ultimately user-guided: analysts use domain knowledge (e.g., known seasonal or interannual cycles) to decide how to cluster elementary components into meaningful signals.

A common issue in SSA is that some elementary components contain contributions from multiple frequencies. This complicates separation when the goal is to isolate, for example, seasonality from interannual variability. The choice of window length is critical here because it defines the trajectory matrix and determines separability. We recommend conducting a sensitivity analysis across a range of window lengths, then minimizing contamination (e.g., by reducing the extent to which seasonality leaks into interannual components or interannual variability contaminates long-term trends).

### Empirical Orthogonal Function Analysis

Empirical Orthogonal Function (EOF) analysis is a data-driven multivariate technique used to decompose spatiotemporal datasets into orthogonal modes of variability. It is mathematically equivalent to applying Principal Component Analysis (PCA) to a spatiotemporal field and is particularly well-suited for identifying dominant spatial patterns and their coherent temporal evolution. The method relies on the eigenvalue decomposition of the dataset’s covariance matrix, yielding a set of spatial modes (EOFs) that capture the maximum variance in the data, along with associated temporal coefficients (principal components) that describe their temporal evolution.

Mathematically, the decomposition follows:

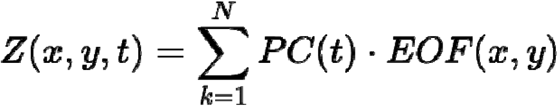

### Model Formulation and Validation

We modeled dengue incidence and altitudinal distribution using Generalized Additive Models (GAMs). To account for the nature of the response variables, we employed a Poisson likelihood with log-link function for case counts. To prioritize structural stability and prevent overfitting, the basis functions for the smooth terms were constrained using 2 knots of degree 2. This conservative parameterization ensures that the identified non-linearities reflect robust, large-scale relationships between climate drivers and disease dynamics rather than local noise.

To assess the predictive robustness of these models and account for temporal dependencies, we employed a forward-chaining (rolling origin) cross-validation scheme. The time series was divided into five sequential folds. We iteratively trained the models on the preceding folds to predict the subsequent “unseen” fold (e.g., training on folds 1–2 to predict fold 3, and eventually training on folds 1–4 to predict fold 5). This approach ensures that the model’s predictive skill is evaluated by its ability to forecast future disease dynamics using historical climate-driven signals.

## Acknowledgements

The authors would like to acknowledge Paloma Trascasa Castro for insightful discussions on the relative robustness of ENSO teleconnections compared to ENSO SST indices in climate change projections. Mauricio Santos-Vega acknowledges the ICTP Associates Program for providing support and opportunities that contributed to this work. Adrià San José acknowledges support from the Severo Ochoa mobility grant at the Barcelona Supercomputing Center, which enabled him to undertake a research stay at Universidad de los Andes. Rachel Lowe acknowledges support from the Wellcome Trust (HARMONIZE 224694/Z/21/Z and IDExtremes 226069/Z/22/Z) and the European Union’s Horizon Europe research and innovation program (E4Warning, grant agreement 101086640, and IDAlert, grant agreement 101057554).

## Supplementary Figures

**Fig. S1.**
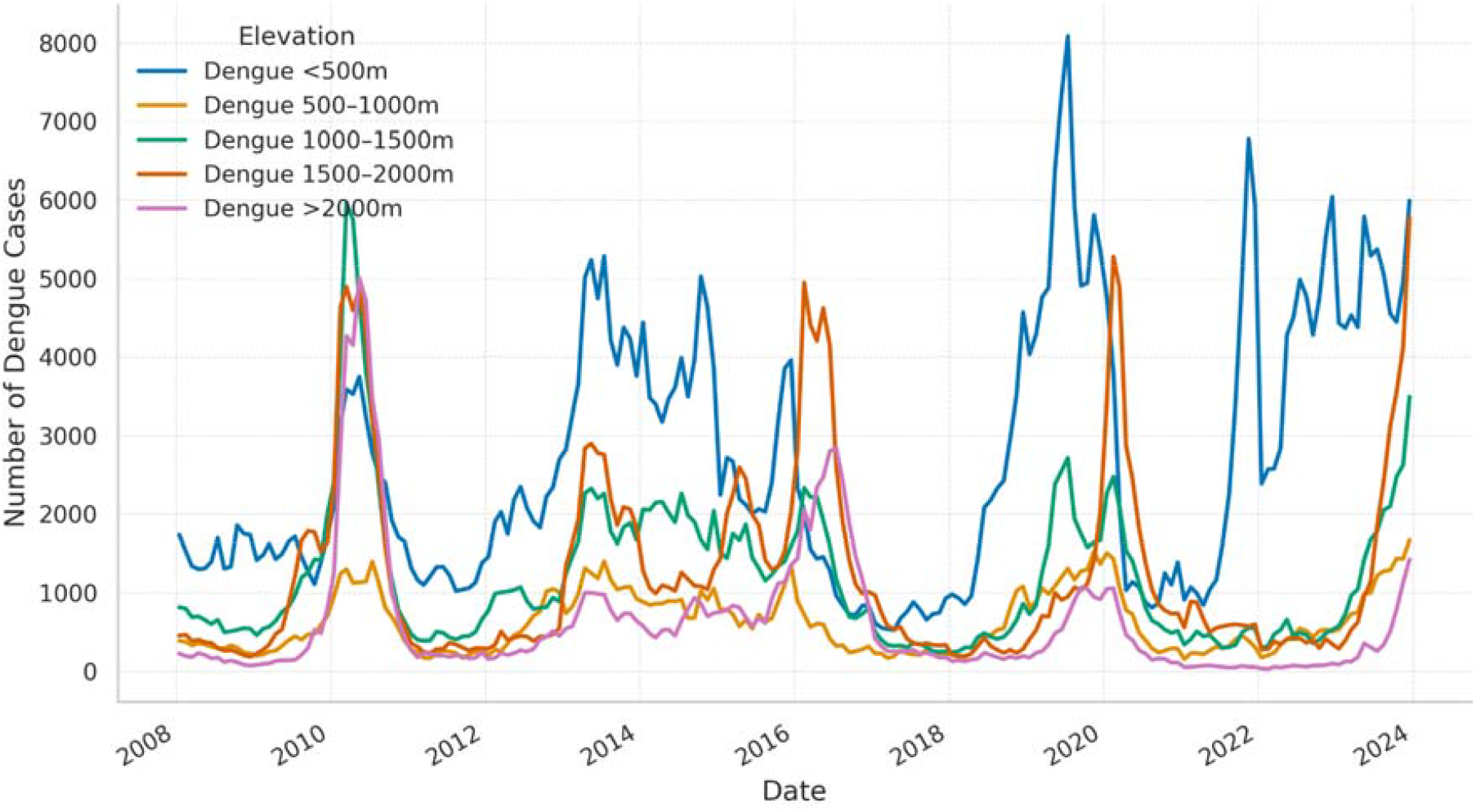
Temporal patterns of dengue cases across elevation bands. Transmission in lowland areas (<500 m) is temporally decoupled from highland transmission. Above 500 m, dengue dynamics are synchronized across elevation ranges.

**Fig. S2.**
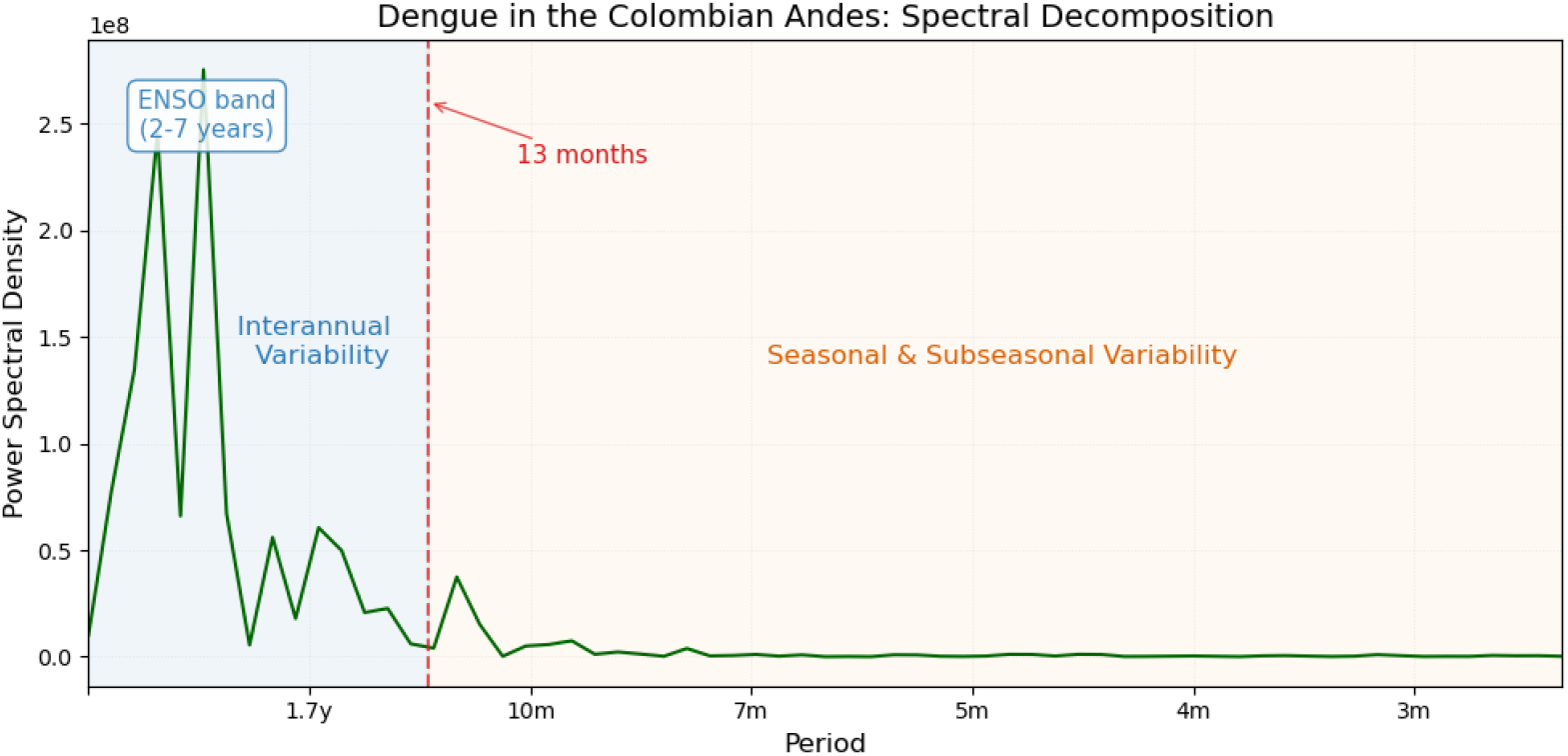
Spectral Decomposition of dengue dynamics in the Colombian Andes, showing most variability is interannual and especially concentrated in the ENSO (2-7 years) band.

**Fig. S3.**
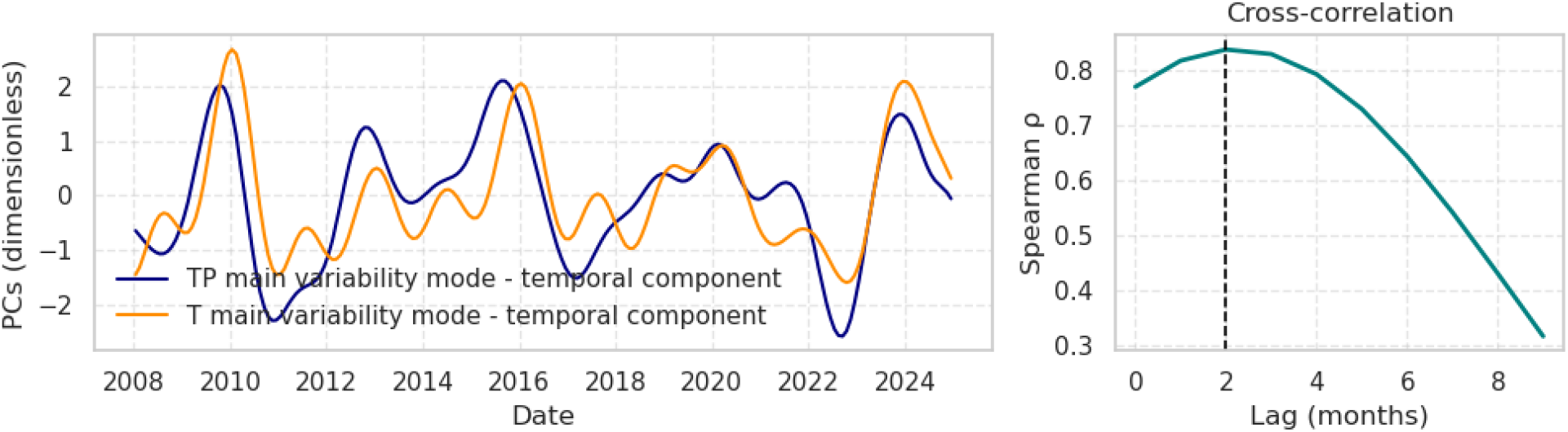
Principal components (PCs) representing the interannual variability of temperature (T) and precipitation (TP) are shown. Cross-correlation analysis reveals a very strong association between the two variables (Spearman’s ρ > 0.85) when precipitation leads temperature by approximately two months.

**Fig. S4.**
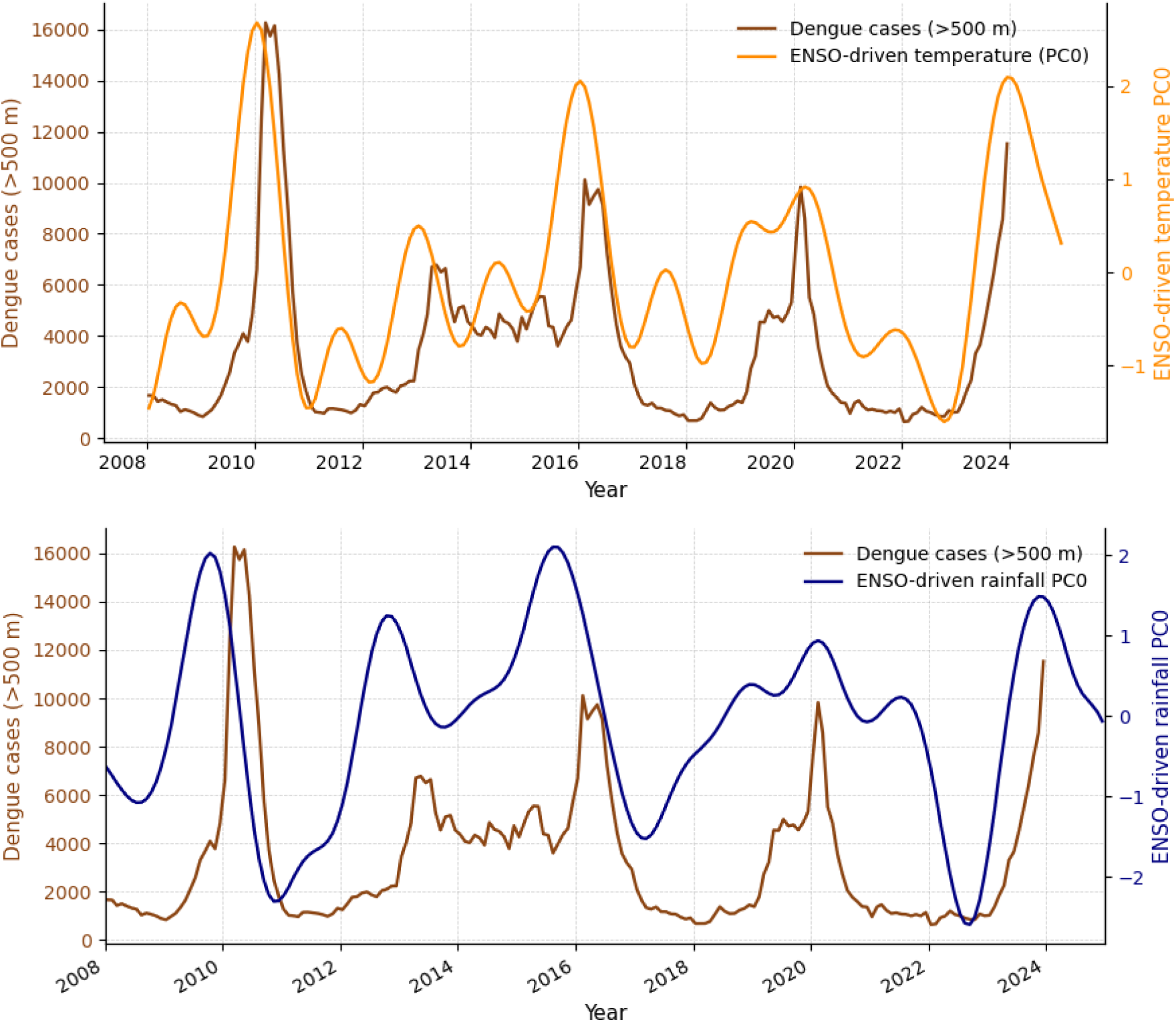
Dengue dynamics at high elevations in relation to ENSO-driven climate variability. Dengue cases occurring above 500m are shown alongside the principal components (PCs) of temperature and precipitation variability associated with ENSO. The combined influence of these climate modes exerts a strong control on high-elevation dengue dynamics.

**Fig. S5.**
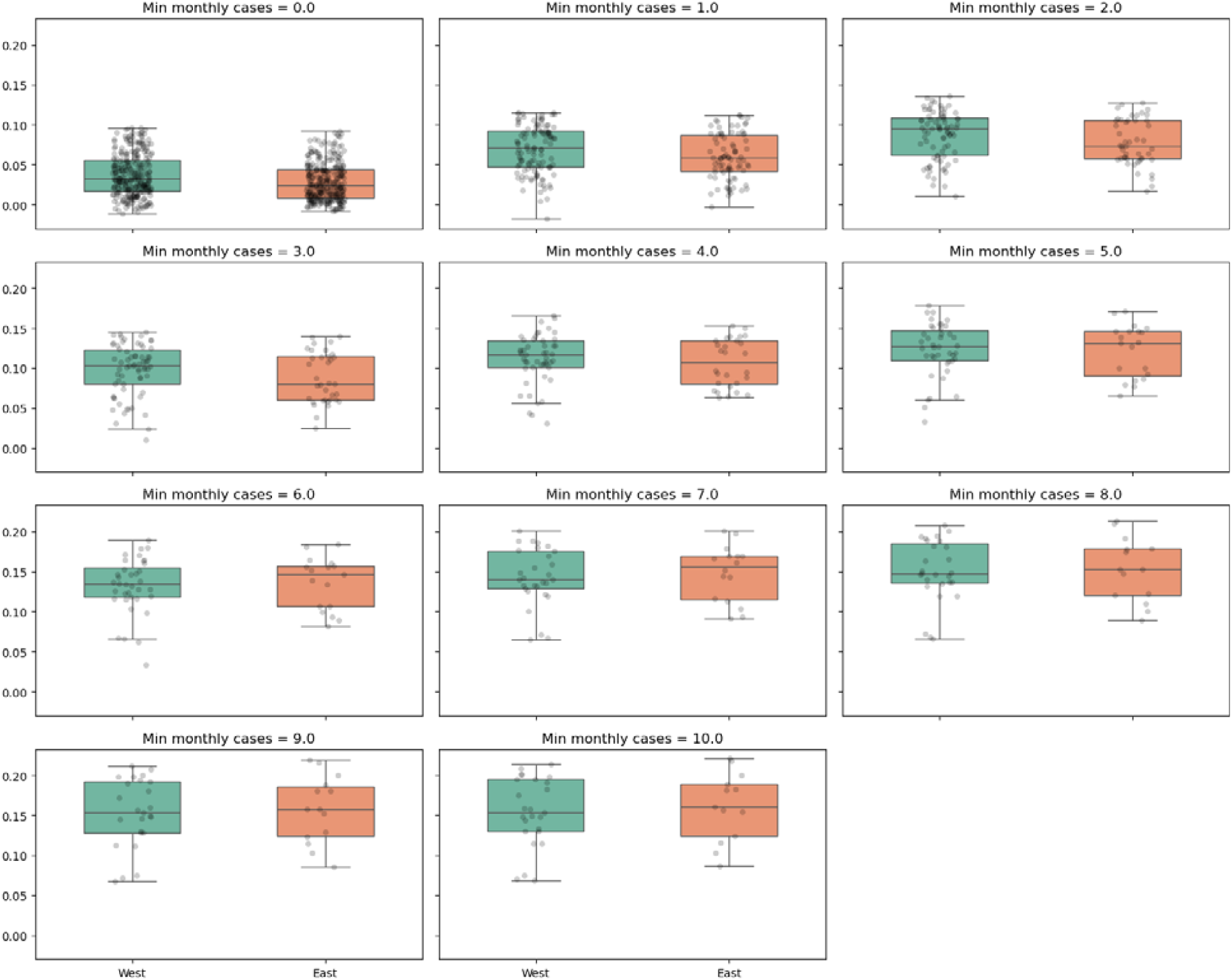
Iterative PCA results stratified by ridge for dengue dynamics at the municipal level. The iterative procedure explores different thresholds for minimum monthly cases to reduce noise and ensure robustness. One ridge (Western) consistently exhibits higher case counts and is therefore more robust to noise, which could otherwise inflate apparent ENSO sensitivity relative to the Eastern ridge. This motivates the use of the iterative thresholding approach, which confirms the same overall sensitivity regardless of the threshold selected.

**Fig S6.**
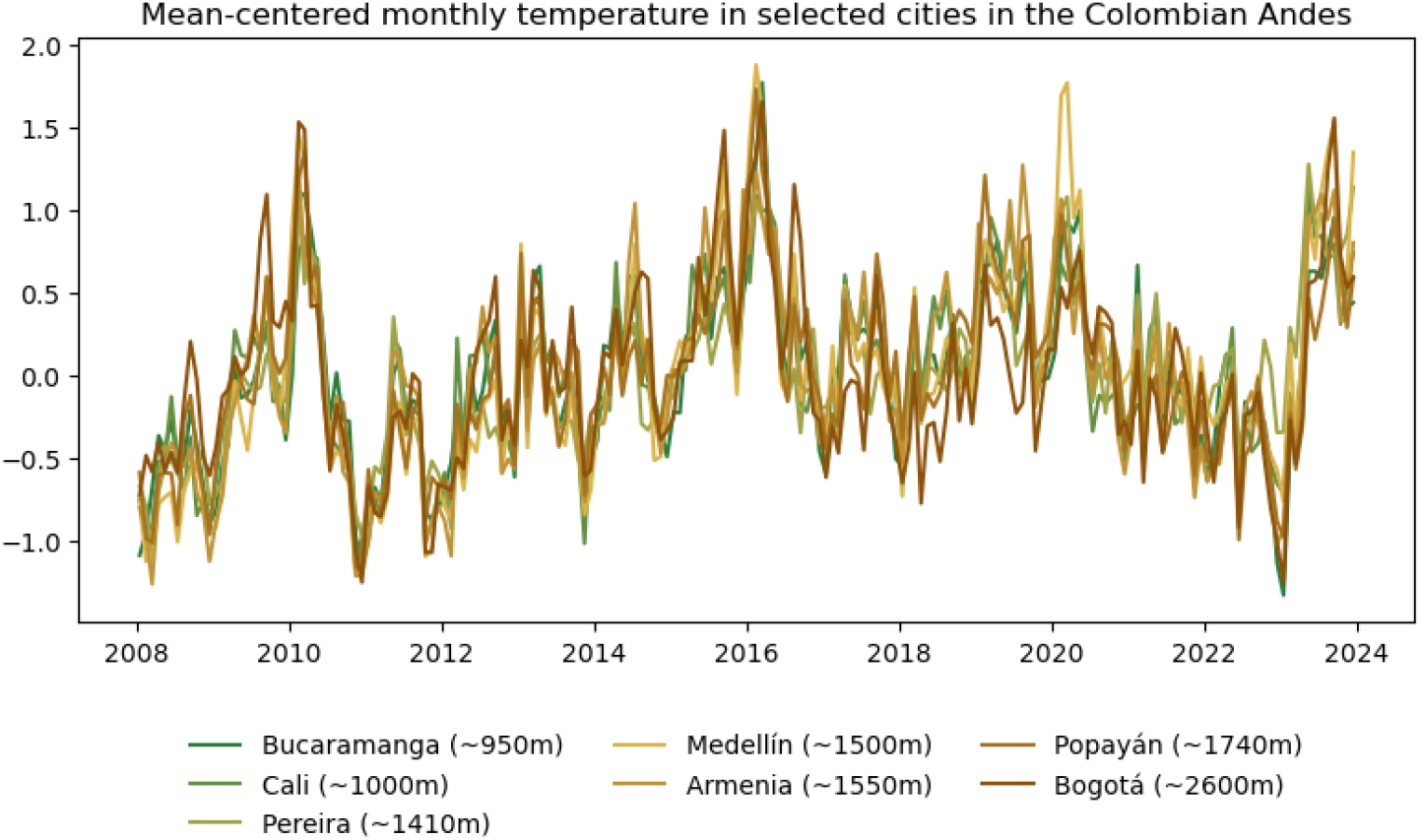
Mean-centered temperature anomalies across Colombian cities spanning the Andean elevational gradient. Colors transition from green (low elevation) to brown (high elevation). While minor differences among cities are visible, no systematic elevational gradient or coherent altitudinal signature emerges in the temperature anomalies themselves.

**Fig. S7.**
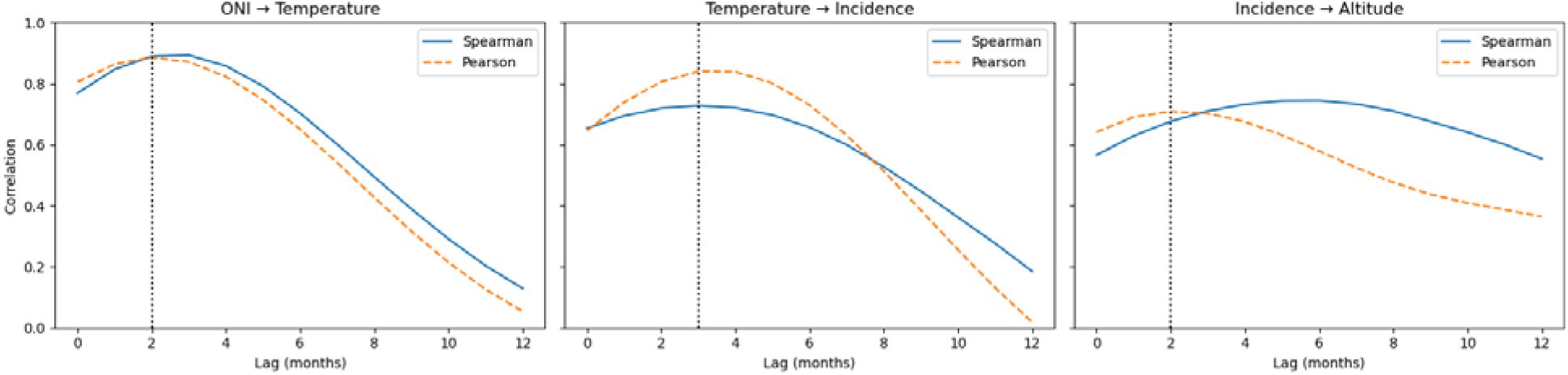
Cross-correlation analysis between ONI and local temperature, local temperature and dengue incidence, and incidence and altitude. Correlations were computed using both Pearson and Spearman coefficients. Pearson correlation, being sensitive to the magnitude of anomalies, more strongly captures the dominant lag associated with ENSO peak events. Spearman correlation assesses rank-order consistency and is therefore more robust to the influence of extreme values, providing a complementary characterization of the monotonic relationship across the full range of variability

**Fig. S8.**
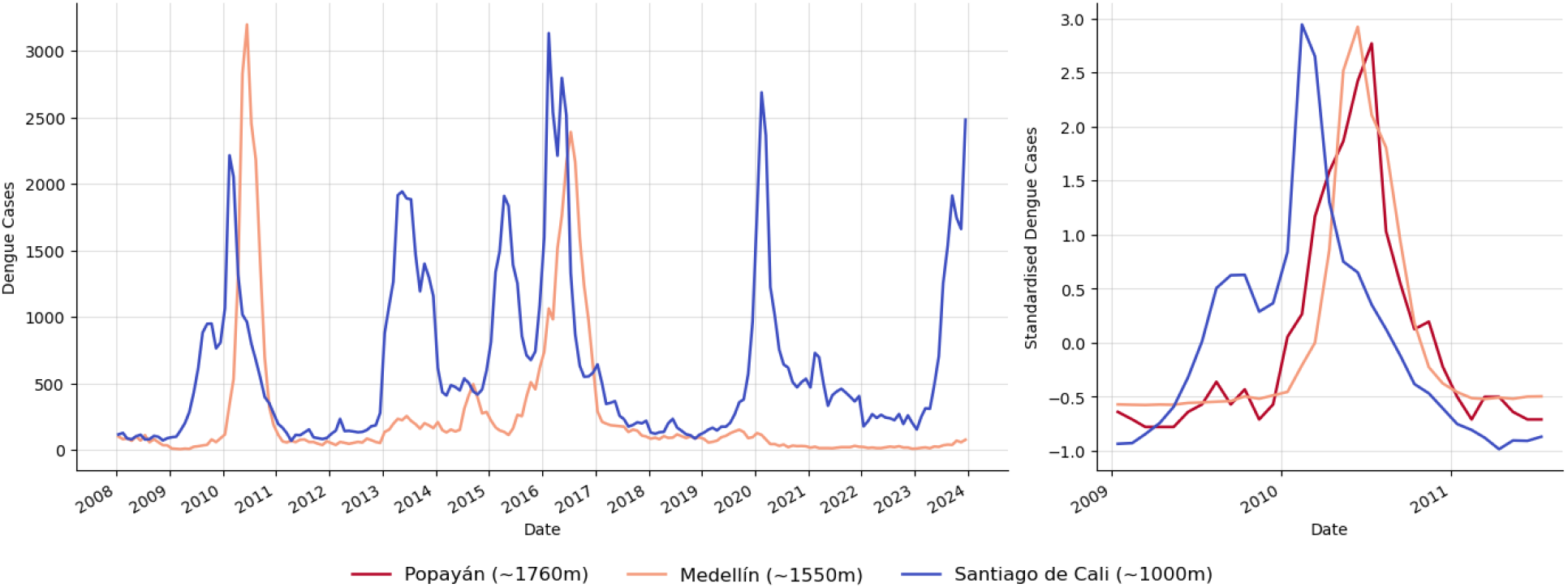
Delayed outbreak onset at increasing elevations. **(A)** Dengue temporal dynamics in Cali (∼1000 m) and Medellín (∼1550 m), the two main dengue hotspots in Colombia. Higher-altitude Medellín exhibits a systematically delayed epidemic response relative to Cali; **(B)** Zoom on the 2010 epidemic, one of the most widespread outbreaks on record in Colombia, extended to include Popayán (∼1,760 m), one of the highest cities to show strong transmission during the peak, to span a broader elevational gradient. Cases are standardized (z-scores) to remove differences in outbreak magnitude. Epidemic onset and peak are progressively delayed with increasing altitude.

**Fig. S9.**
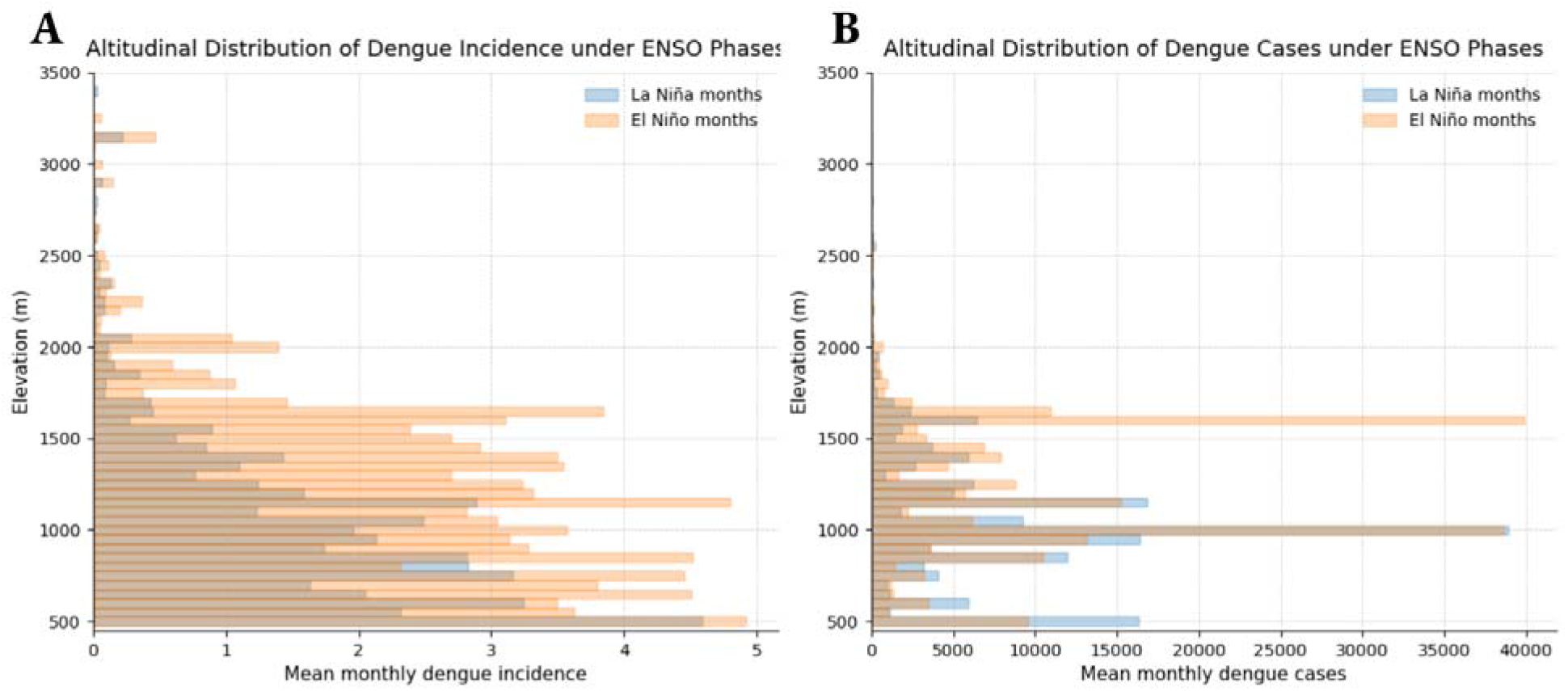
Altitudinal distribution of dengue incidence **(A)** and cases **(B)** across Colombia. The 2300□m threshold applied in Fig. 2A is removed. Very few cases occur above 2000□m, as illustrated in (B). Certain sparsely populated elevation bands around ∼3200□m artificially elevate incidence in (A), but these values are not meaningful, as are extremely noisy.

**Fig. S10.**
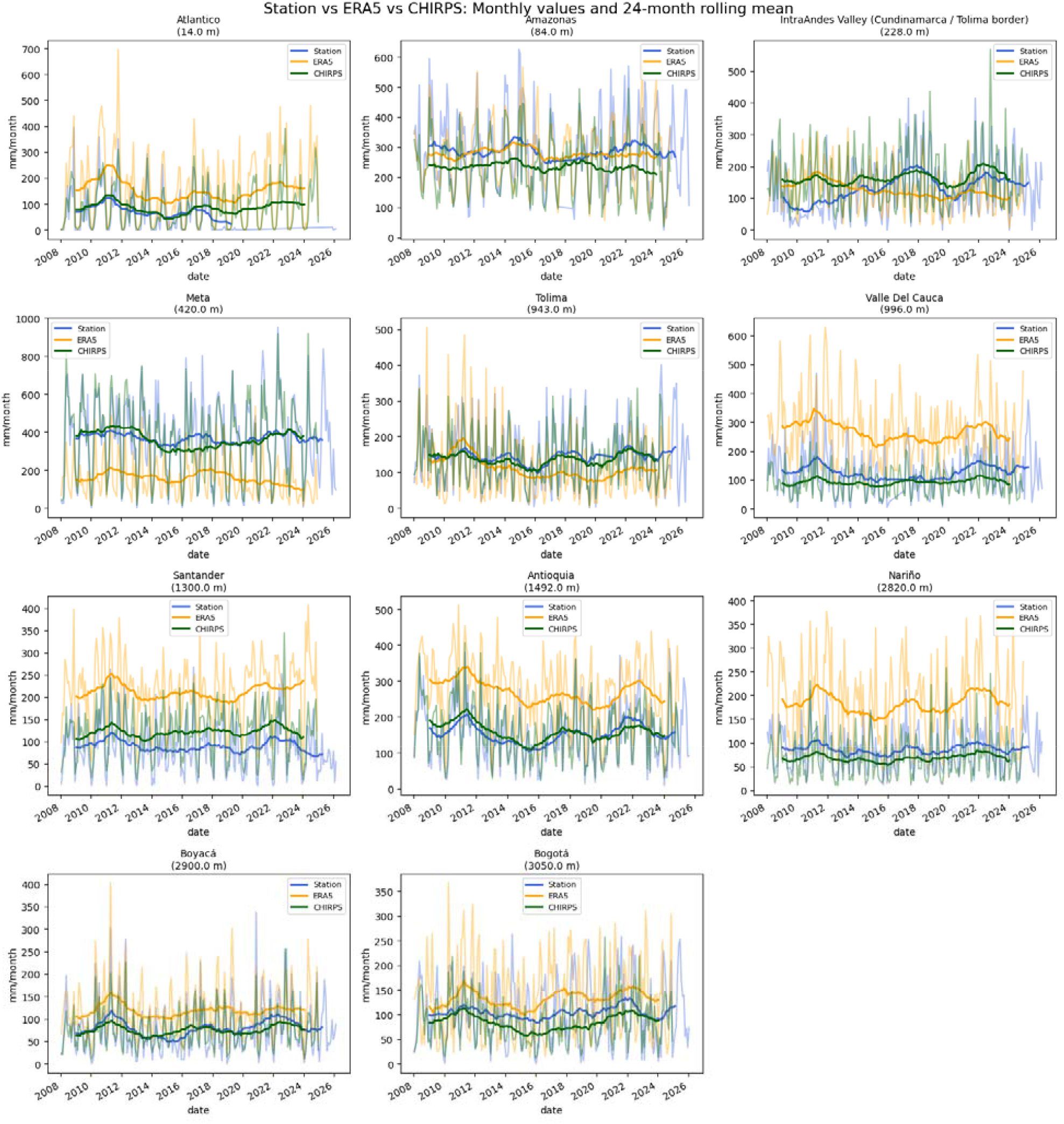
Validation of low-frequency precipitation variability against station observations. Interannual rainfall anomalies derived from ERA5 and CHIRPS are compared with station records across representative locations spanning the main climatic regions of Colombia: cities along the western and eastern Andean ridges, the inter-Andean Magdalena Valley (Cundinamarca/Tolima border), the Caribbean coast (Atlántico), and the Amazon basin (Amazonas).

**Fig. S11.**
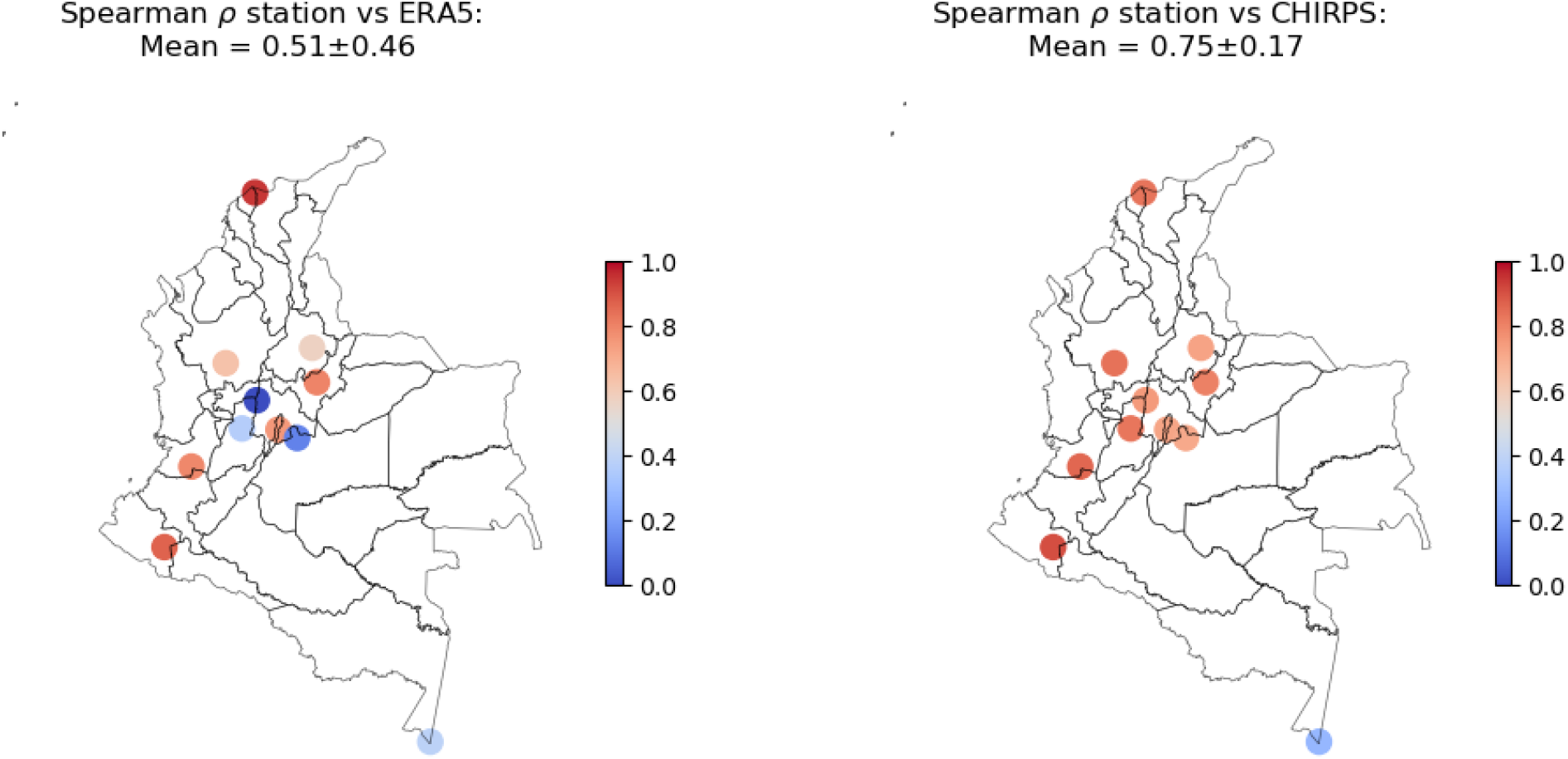
Spearman rank correlations between interannual precipitation anomalies from ERA5 and CHIRPS and station observations across Colombia. Each station is shown at its geographic location, with marker color indicating the strength of agreement with each gridded product. The spatial distribution of stations spans the main climatic and topographic regions of the country, allowing assessment of dataset performance across contrasting environments, but especially in the Andes. CHIRPS shows better agreement with station data.

## APPENDIX A

We estimated relative vectorial capacity following the standard formulation of (Mordecai et al, 2019), parameterized for the *Ae. aegypti-*DENV system:

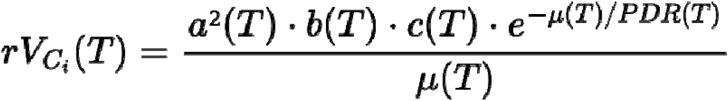

Each temperature-dependent parameter is described by either a Brière or quadratic function.

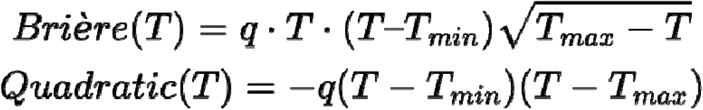

The parameters and their coefficients are as follows:

□ a(T) denotes the Biting Rate and is described by a Brière function with q=2.02, Tmin = 13.8 and Tmax = 40.0
□ b(T) denotes the transmission probability (mosquito to host) and is described by a Brière function with q=8.33·10^−4^, Tmin =17.2 and Tmax =35.8
□ c(T) denotes the infection probability (host to mosquito) and is described by a Brière function with q=4.88·10^−4^, Tmin=12.7 and Tmax=37.4
□ PDR stands for the Parasite development rate and is described by a Brière function with q=6.13·10^−5^ and Tmin= 10.3 and Tmax=45.6
□ Mu stands for the mortality rate. It’s the inverse of the lifespan, which can be described with a quadratic formula with q=1.44·10^−1^, Tmin=9.0 and Tmax=37.7

The complete formulation of vectorial capacity includes *m*, the mosquito-to-human ratio. Because we standardized the elevational centroid of vectorial capacity to focus on its temporal dynamics rather than its absolute baseline altitude, the absolute value of *m* does not affect our primary inference. Furthermore, even if *m* were dynamic, increasing during warm or wet periods and declining during cold or dry ones, this would not alter our conclusions, as the adjacent urban centers considered here experience broadly similar seasonal and interannual climate patterns across the Colombian Andes.

## Notes

### Competing Interest Statement

The authors have declared no competing interest.

### Summary of Updates

Expanded revision incorporating additional mechanistic detail and clarifications

